# *ChemXTree*:A Tree-enhanced Classification Approach to Small-molecule Drug Discovery

**DOI:** 10.1101/2023.11.28.568989

**Authors:** Yuzhi Xu, Xinxin Liu, Jiankai Ge, Wei Xia, Cheng-Wei Ju, Haiping Zhang, John Z.H. Zhang

## Abstract

The rapid advancement of machine learning, particularly deep learning, has propelled significant strides in drug discovery, offering novel methodologies for molecular property prediction. However, despite these advancements, existing approaches often face challenges in effectively extracting and selecting relevant features from molecular data, which is crucial for accurate predictions. Our work introduces ChemXTree, a novel graph-based model that integrates tree-based algorithms to address these challenges. By incorporating a Gate Modulation Feature Unit (GMFU) for refined feature selection and a differentiable decision tree in the output layer. Extensive evaluations on benchmark datasets, including MoleculeNet and eight additional drug databases, have demonstrated ChemXTree’s superior performance, particularly in feature optimization. Permutation experiments and ablation studies further validate the effectiveness of GMFU, positioning ChemXTree as a significant advancement in molecular informatics, capable of rivaling state-of-the-art models.

## 1 Introduction

In recent decades, the development of machine learning has remarkably accelerated the pace of drug discovery. ^1,2^ This acceleration is primarily because machine learning, unlike traditional drug discovery methods, can quickly process large datasets to identify crucial ADMET (Absorption, Distribution, Metabolism, Excretion, and Toxicity) properties of drug compounds. ^3–5^ By efficiently predicting factors such as distribution coefficients, free energy, solubility, stability, e.g., machine learning enhances the efficacy and safety assessment of potential drugs. ^6,7^ As one of the most rapidly evolving subsets of machine learning, deep learning has made significant strides in molecular property prediction. In molecular property prediction, deep learning offers the advantage of avoiding the need for pre-defined features, a limitation in traditional machine learning approaches. By providing an end-to-end mapping from input to output, deep learning categorizes its primary methodologies for molecular feature extraction based on distinct computational frameworks: models utilizing visual grid encoding often rely on Convolutional Neural Networks (CNNs) to capture spatial molecular features, ^8,9^ sequence-based models employ architectures like Recurrent Neural Networks (RNNs) and Transformers to leverage molecular notations, ^10,11^ and graph-based models harness the power of Graph Neural Networks(GNN) to model the intricate topological relationships between atoms and bonds in molecules. ^12,13^ Currently, much of the deep learning in molecular property prediction is dedicated to enhancing the representation capabilities of molecules, aiming to extract more information directly from molecular structures. Several notable approaches have demonstrated superior feature extraction compared to traditional molecular fingerprints, resulting in tangible benefits during training. ^14,15^ However,in recent work, deep learning-based methods have not consistently outperformed methods employing decision trees. ^16,17^ In molecular representation, the overall process can be regarded as two main parts: feature extraction and output prediction. Most people opt to feed the extracted features directly into a Feed-Forward Neural Network (FFN) without any additional feature engineering or selection because deep learning can automatically highlight the most important features and downplay the less relevant ones. Nevertheless, specific substructures and atomic attributes are crucial in predicting molecular characteristics. Thus, additional feature weight optimization could boost the model’s performance. Deng et al. proposed XGraphBoost which substituted Extreme Gradient Boosting(XGBoost) for the FFN component in the D-MPNN model. ^18,19^ Their comparative experiments demonstrate that this modification enhances the model’s performance. This improvement is attributed to the inherent ability of tree-based models to automate feature selection.

To improve model performance, tree-based techniques have been investigated in molecular property prediction. Liu et al. introduced an N-Gram model and employed Random Forest(RF) and XGBoost as output layers, replacing the traditional linear network. ^20^ These tree-based methods are adept at leveraging intricate embedding information from complex models to boost performance. Although tree-based output layers have found numerous applications in molecular prediction, most existing research primarily focuses on the integration of traditional decision trees into machine learning frameworks. ^21–23^ In contrast, there is a notable scarcity of literature that explores the incorporation of neural decision trees with backpropagation algorithms as part of a unified model architecture for molecular prediction. Zhan et al. recently introduced the Graph Neural Tree. This innovative model employs a Learning Binary Neural Tree (LBNT) in the output layer. ^24^ In four cases, LBNT outperforms both the FFN output commonly used in Graph Neural Networks and the output generated by RF.

In this study, we propose ChemXTree, a tree-enhanced graph representation model for molecular property prediction. To boost the feature selection capability, we introduce a new module named Gate Modulation Feature Unit (GMFU) to refine and select the most informative features. After the feature selection process, a differentiable decision tree is further incorporated as the prediction output layer. We conducted comprehensive evaluations using standard benchmark datasets from the MoleculeNet and 8 additional drug databases. ^25^ The results demonstrated that ChemXTree exhibits significant competitiveness. Additionally, we executed permutation experiments on the output layer to evaluate the effect on the overall model performance. We also performed ablation studies on the GMFU and developed a LSTM-based variant of GMFU. In general, we demonstrate that, even without pre-training, the combination of GMFU and decision trees to enhance feature selection can rival the performance of existing SOTA pre-trained models in classification tasks.

## 2 Result and Discussion

### 2.1 ChemXTree Workflow

ChemXTree is specifically designed to address classification problems in small drug molecules.To achieve this, the architecture of ChemXTree leverages GNN for molecular encoding and encoded output is then fed into an innovative module, dubbed GMFU, dedicated to optimized feature selection. After that, ChemXTree employs a Differential Decision Tree, enhancing the model’s classification capabilities.

To more specific, as depicted in Figure 1, ChemXTree initiates its process by first transforming SMILES of molecules into graphs. In these graphs, atoms, along with multi-scale information such as charge and valency, serves as vertices(nodes), while chemical bonds act as edges. Subsequently, in the molecular representation learning stage, these initially encoded molecules are passed to Message Passing Neural Network (MPNN) for further processing and refinement of the molecular encoding. ^26^ Upon completion of the encoding training, the MPNN equipped with trained weights then functions as an encoder, encoding the validation and test sets, respectively. With this method, the encoded representations of the molecules in train/validation/test sets are obtained. To enhance the robustness of molecular classification, we employ an ensemble approach. Specifically, the final molecular encoding is obtained by summing the outputs of five independently trained models for each molecule.

**Figure 1:**
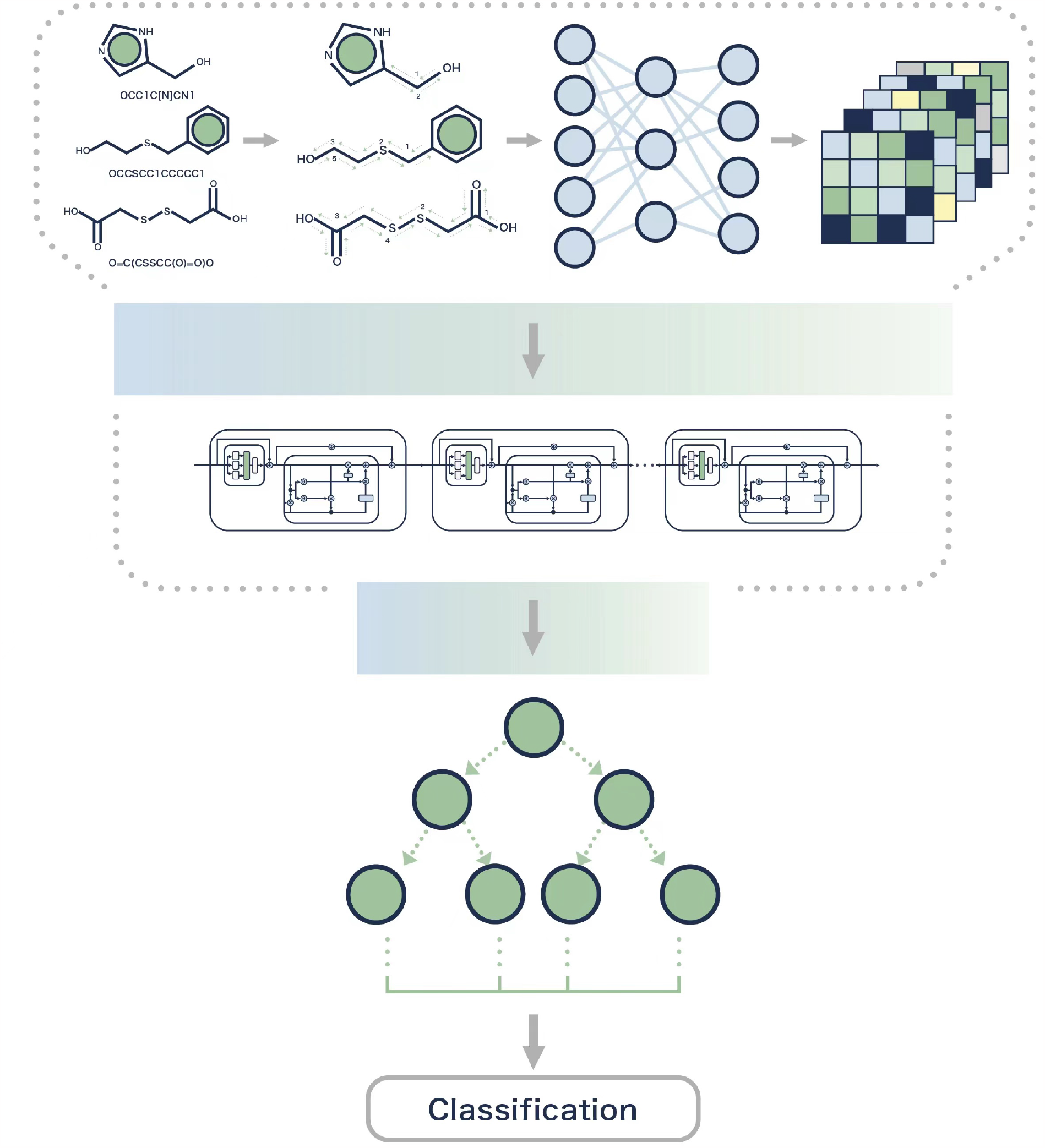
The workflow of ChemXTree: a strategy specifically engineered for the classification of small drug molecules, where molecules are converted into graph-based representations and then encoded through an MPNN module. The encoded outputs from MPNN module are augmented fivefold by an ensemble of models, followed by feature selection via GMFU modules, culminating in classification through a Differential Decision Tree.

With the encoded molecular representations, ChemXTree aims to intensify the identification of task-relevant features for downstream tasks. To achieve this, we employ a network architecture composed of a series of GMFUs. Inspired by Gated Recurrent Units (GRU) and other variants like Gated Adaptive Network using similar gated mechanism, GMFU is designed for further feature selection.^27–29^ Specifically, each GMFU start with self-attention on the input features, which are then fed into a gated structure equipped with reset and update gates to generate candidate feature representations. The final output of the GMFU is computed by combining the previous hidden states with the current candidate features based on the update gate. Multiple GMFUs can be cascaded in a hierarchical fashion, where the output of one unit serves as the input for the subsequent unit. Different layers of GMFU engage in distinct feature selections, enabling the model to capture molecular representations at varying levels of abstraction. This facilitates the integration of both global and local information into the final feature representation.

These features are subsequently fed into a differentiable decision tree for further employment. To enable continuous optimization and backpropagation of gradients, our decision tree employs soft decisions based on output probabilities, as opposed to traditional hard decisions that only yield 0 or 1. By leveraging an ensemble of different differentiable trees and their weighted outputs, a binary prediction result is generated. For more information about the architecture of the ChemXTree, please see the Materials and Methods section.

### 2.2 Benchmark in MoleculeNet

To comprehensively evaluate the ChemXTree, we utilized the MoleculeNet databases developed by Wu et al. ^25^ ChemXTree is primarily designed for single-target binary classification and therefore, given the computational expense and deployment challenges of multi-task binary classification, coupled with our computational resource constraints, we did not test on tasks with multi-task binary classifications. In this evaluation part, we adhered to the data processing and evaluation method presented in previous work. After obtaining the optimal hyperparameters through Bayesian optimization and running the model three times, we conducted a comparison with various existing models.

As shown in Table 1, ChemXTree performs comparably or slightly better than the state-of-the-art (SOTA) models in all the datasets. Notably, ChemXTree excels in predicting tasks like BBBP, BACE, and ClinTox. A potential factor could be ChemXTree’s strategic feature selection combined with its tree-based output layer, offering an equilibrium between fitting the data and generalizing well, especially with small datasets. Specifically, compared to D-MPNN ^26^ where similar molecular feature extraction is applied, ChemXTree demonstrates a consistent trend of improvement across all datasets, boosting the ROC-AUC by 6% for both BBBP and BACE tasks. This indicates that enhancing feature selection can sighnificanty boost the simple GNN structures. Moreover, models such as N-GramRF, ^20^ N-GramXGB, PretrainGNN, ^30^ GROVERlarge, ^31^ GROVERbase, MolCLR, ^32^ and Uni-Mol ^33^ undergo pre-training and fine-tuning process. In contrast, ChemXTree attains comparable results without this step, indicating the potential of its structure and algorithms even in the absence of pre-training.

**Table 1:** Performance of ChemXTree and Other Models Across BBBP, BACE, HIV, and ClinTox Datasets with Best ROC-AUC Score Denoted in Bold.

| Datasets | BBBP | BACE | HIV | ClinTox |
| --- | --- | --- | --- | --- |
| D-MPNN | 71.0 | 80.9 | 77.1 | 90.6 |
| Attentive FP<sup>34</sup> | 64.3 | 78.4 | 75.7 | 84.7 |
| N-GramRF | 69.7 | 77.9 | 77.2 | 77.5 |
| N-GramXGB | 69.1 | 79.1 | 78.7 | 87.5 |
| PretrainGNN | 68.7 | 84.5 | 79.9 | 72.6 |
| GROVERbase | 70.0 | 82.6 | 62.5 | 81.2 |
| GROVERlarge | 69.5 | 81.0 | 68.2 | 76.2 |
| GraphMVP | 72.4 | 81.2 | 77.0 | 79.1 |
| MolCLR | 72.2 | 82.4 | 78.1 | 91.2 |
| GEM<sup>35</sup> | 72.4 | 85.6 | 80.6 | 90.1 |
| Uni-Mol | 72.9 | 85.7 | **80.8** | 91.9 |
| ChemXTree(Our) | **75.6** | **86.2** | 80.6 | **92.3** |

### 2.3 Benchmark in Drug Datasets

For a comprehensive benchmark, we tested ChemXTree on 8 additional drug datasets, including AMES, CYP2C9_Substrate, CYP2D6_Substrate, CYP3A4_Substrate, CYP2C9_Inhibition, CYP2D6_Inhibition, CYP3A4_Inhibition, and Bioavailability. These datasets have been widely adopted in previous works as benchmarks. ^40^ Specifically, our dataset selection considers AMES, CYP2C9_Inhibition, CYP2D6_Inhibition and CYP3A4_Inhibition as large-scale datasets, and CYP2C9_Substrate, CYP2D6_Substrate, CYP3A4_Substrate and Bioavailability as small-scale datasets, to cover diverse data characteristics and model capabilities. We compare our ChemXTree against 10 SOTA methods, spanning from conventional machine learning algorithms e.g. XGBoost to emerging deep graph neural networks including DMPNN, AttentiveFP, Graph Attention Network(GAT) et all, as well as pre-trained models like Uni-Mol and GROVER (GROVERbase is chosen as the representative of GROVER due to their performance in MoleculNet) .The detailed benchmark model information could be found in the Materials and Methods. For comparison, we optimize hyperparameters and follow the same dataset splitting and evaluation metrics(ROC-AUC) as MoleculeNet recommend.

As shown in Table 2, our ChemXTree achieves superior average ROC-AUC scores across all 8 datasets compared to other methods, demonstrating its strong generalization capability. Specifically, ChemXTree attains the best ROC-AUC performance on the AMES, CYP2C9, CYP2D6, CYP3A4, and Bioavailability datasets. On the larger CYP2C9_Inhibition, CYP2D6_Inhibition and CYP3A4_Inhibition datasets, the pre-trained Uni-Mol model secures the best ROC-AUC, while ChemXTree and GROVER achieved second-ranked results. This is reasonable since Uni-Mol benefits from pre-training on external millions of molecular data, granting better generalization to represent unseen cases. In contrast, ChemXTree learns representations from scratch solely based on the training data, yet still surpasses other methods without pre-training, verifying its modeling effectiveness. Taken together, these benchmark experiments validate ChemXTree’s superiority and competitiveness on molecular property prediction tasks in both small and large-scale datasets.

**Table 2:** Comparative Analysis of ROC-AUC Scores Across Eight Drug Datasets for ChemXTree and Other Leading Models in AMES, CYP Enzyme Substrate and Inhibition, and Bioavailability, with Top Performances in Bold and Second Best Underlined.

| Datasets | AMES | CYP2C9_Substrate | CYP2C9_Inhibition | CYP2D6_Substrate | CYP2D6_Inhibition | CYP3A4_Substrate | CYP3A4_Inhibition | Bioavailability | Average |
| --- | --- | --- | --- | --- | --- | --- | --- | --- | --- |
| DMPNN | 86.9 | 60.0 | 87.2 | 70.0 | 84.0 | 59.9 | 85.9 | 68.4 | 75.3 |
| Attentive FP<sup>34</sup> | 87.4 | 67.0 | 86.6 | 70.0 | 83.3 | 60.9 | 85.7 | 72.4 | 76.7 |
| XGBoost | 82.7 | 51.0 | 81.1 | 67.1 | 79.2 | 62.5 | 81.3 | 58.9 | 70.5 |
| GAT<sup>36</sup> | 87.2 | 67.7 | 83.5 | 70.7 | 83.5 | 60.0 | 84.6 | 69.4 | 75.8 |
| MAT<sup>37</sup> | 83.6 | 60.6 | 78.1 | 71.3 | 80.0 | 59.7 | 79.6 | 61.5 | 71.8 |
| GROVER | 83.0 | 69.3 | 85.8 | 70.3 | 81.2 | 69.5 | 88.7 | 73.7 | 77.9 |
| InfoGraph<sup>38</sup> | 86.7 | 57.6 | 87.2 | 77.5 | 81.3 | 56.7 | 82.7 | 66.5 | 74.5 |
| AutoML | 89.0 | 61.1 | 89.1 | 64.0 | 86.7 | 65.5 | 88.1 | 62.7 | 75.8 |
| Graphormer<sup>39</sup> | 77.5 | 43.1 | 80.7 | 60.1 | 76.3 | 68.9 | 75.0 | 54.9 | 67.1 |
| Uni-Mol | 88.4 | 62.9 | **91.0** | 75.0 | **88.2** | 60.6 | **89.0** | 72.0 | <u>78.4</u> |
| ChemXTree(our) | **89.1** | **69.5** | <u>89.2</u> | **78.4** | <u>86.9</u> | **69.6** | 88.2 | **77.2** | **81.0** |

### 2.4 Substitution and Ablation Experiments

To analyze ChemXTree’s enhancements, we took the molecular encoding from the top-performing ChemXtree on BACE/BBBP/HIV datasets to conduct the ablation experiments by swapping its prediction output with a simple FFN and Xgboost. The FFN configuration was from Chemprop’s recommended default of 300 hidden layers and was trained over 1000 epochs. We conducted a grid search over the following hyperparameters: batch sizes of {16, 32, 64} and learning rates of {0.001, 0.002, 0.004}. For the Xgboost, we employed Bayesian optimization within a defined parameter space: learning rate between 0.004 and 1.0, `max_depth` in the interval [4, 20], and `n_estimators` spanning [20, 400]. We incremented the Bayesian search iterations, starting from 30 and gradually increasing to 50, 200, 300. Besides, the lamda and alpha, which is regularization terms, are in the range of [0,10]. For these two layer substitutions, we recorded the best performance model for comprehensive analysis, respectively.

In Figure 2, the comparison of three output layers using identical encoded inputs shows that ChemXTree outperforms both FFN and Xgboost in terms of classification efficacy. We computed the average ROC-AUC score of each output layers across three datasets, with the performance scores ranking as follows: ChemXTree > Xgboost > FFN. Specifically for the HIV dataset(Figure 2C), which has a significant class imbalance with approximately 1:27 ratio of positives to negatives, tree-based approaches clearly outperform FFN. This can be attributed to the tree methods’ branching mechanisms, which are highly effective at capturing nonlinear relationships within the data. Besides, While XGBoost does not surpass the FFN on the BBBP dataset, its respectable performance in other cases supports its viability as an effective substitute for the FFN layer in a range of computational scenarios.

**Figure 2:**
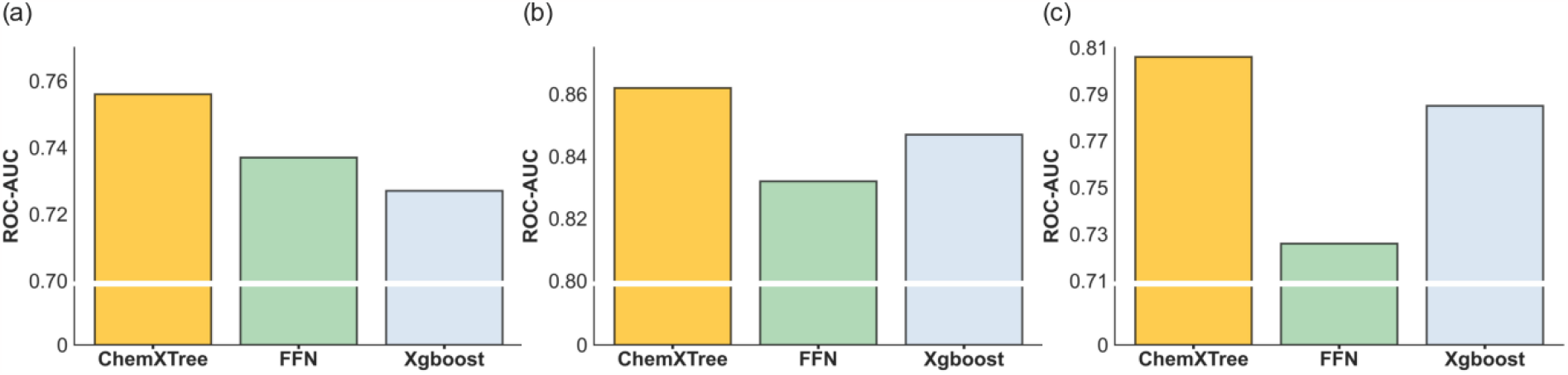
ROC-AUC performance comparison of output layers from ChemXTree, FFN, and Xgboost across BACE, BBBP, and HIV datasets, using uniform encoded inputs

In our exploration of output layers, tree-based methods generally demonstrated superior performance. However, the specific contribution of GMFU was not clearly demonstrated. To address this, we conducted an ablation study to evaluate the effectiveness of GMFU. Employing identical inputs and optimal performance parameters, we compared ChemXTree and ChemXTree variants without GMFU across three datasets. In the figure 3 presented, ‘w/o GMFU’ signifies the ChemXTree configuration excluding its GMFU module. The comparison indicates a decline in model’s performance across three datasets upon removal of the GMFU component. It indicates that the GMFU is integral for optimizing the ChemXTree’s performance due to its enhancement of feature selection.

**Figure 3:**
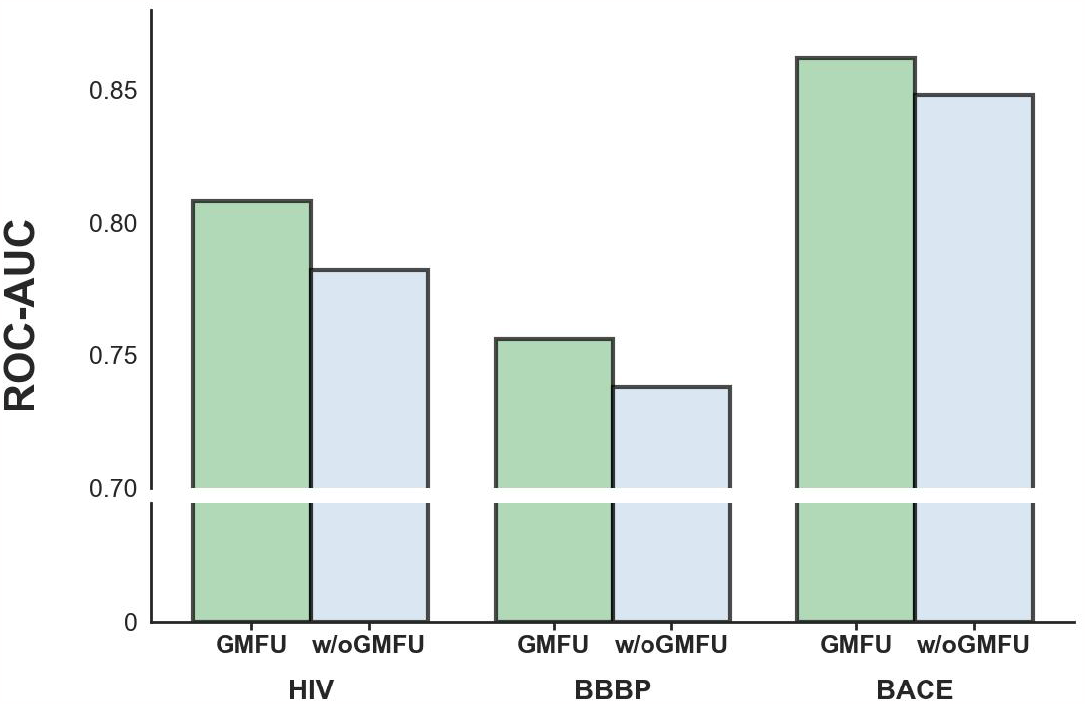
Performance impact of the GMFU module in ChemXTree, comparing the standard model with a variant lacking the GMFU (‘w/o GMFU’) across BACE, BBBP, and HIV datasets

### 2.5 GMFU vs LSTM-based

In addition to the GMFU module, we have also developed a module inspired by Long Short-Term Memory (LSTM) architecture. ^41^ To clearly distinguish between these two modules by mechanism, we designate them as GMFU and LSTM-based respectively. It’s noteworthy that the model adaptation for the LSTM-based variant is nearly identical to the treatment of GMFU. Continuing with the experimental approach delineated in the preceding section, we subjected the BBBP, BACE, and HIV datasets to a comprehensive comparative analysis. As depicted in Figure 4, the original ChemXTree outperforms its LSTM-based variant on all three datasets, achieving a performance boost of 2%.

**Figure 4:**
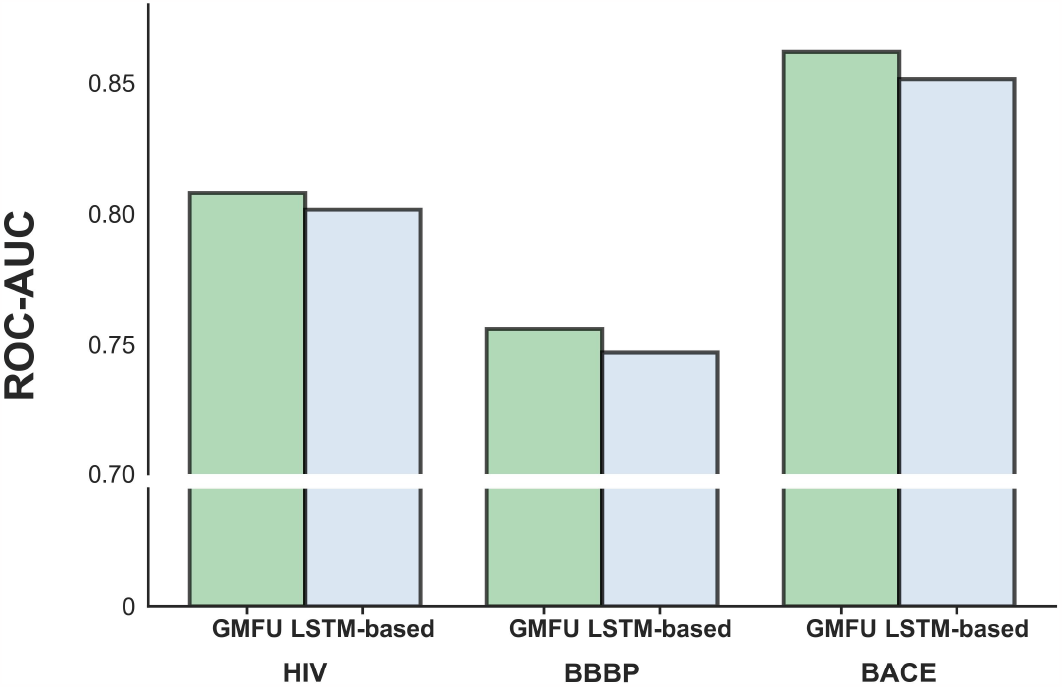
Performance comparison of the original ChemXTree and its LSTM-based variant on the BBBP, BACE, and HIV datasets

**Figure 5:**
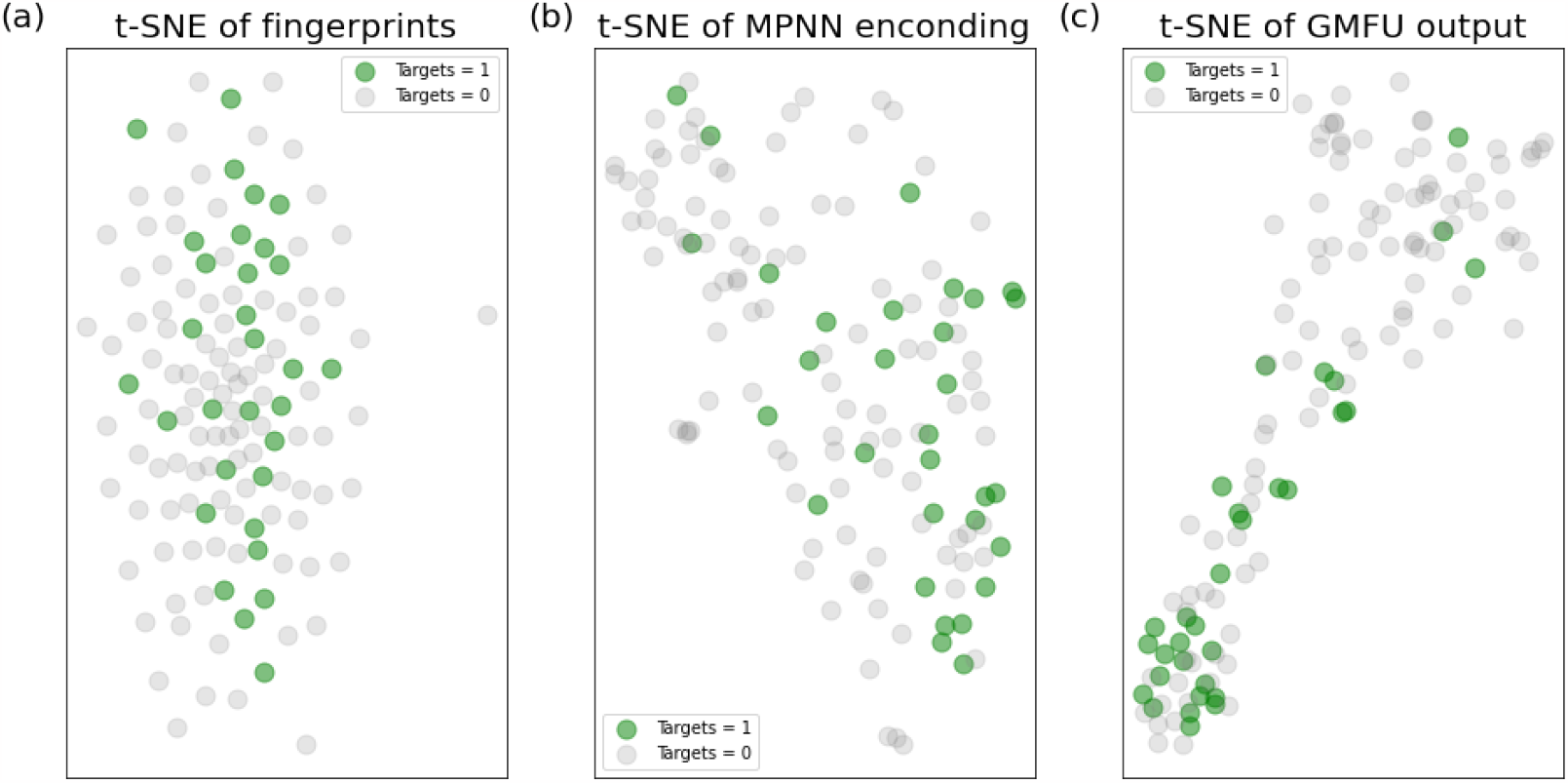
t-SNE visualization of CYP2D6_Substrate testset, showing 2048-bits Morgan Fingerprints, MPNN-encoded Embeddings, and ChemXTree hidden layer Representations, with ground truth ‘1’ and ‘0’ color-coded in green and grey

**Figure 6:**
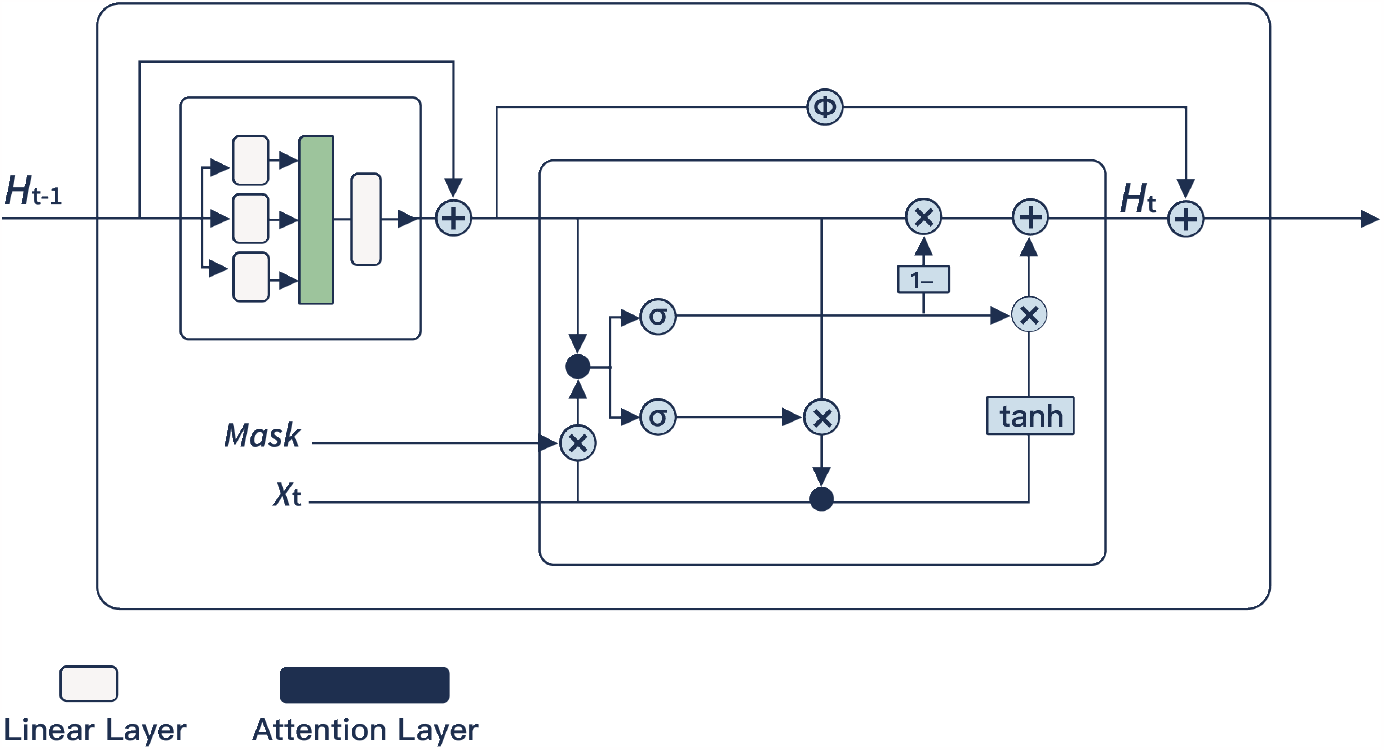
Overview of the GMFU Architecture: Inspired by GRU and GANDALF, GMFU integrates attention mechanisms with gating and residual components.

In contrast to the LSTM-based approach, which employs a complex three-gate architecture for feature selection, the GMFU strategy adopts a more economical two-gate design comprising only reset and update gates. This simplified configuration yields not only a model with fewer parameters, but also a more memory-efficient structure. As evidenced in Table S2, GMFU achieved faster average training times.

### 2.6 Visualization of Model

We use CYP2D6_Substrate testset for t-SNE analysis to visually compare three different representations: 2048-bit molecular Morgan fingerprints, ^42^ MPNN-encoded embeddings, and hidden layer representations from the ChemXTree model before feeding into differential decision tree. ^43^ The plots in t-SNE were generated where ground truth labels ‘1’ and ‘0’ were denoted in green and grey, respectively. The t-SNE results revealed dispersed distributions for the Morgan fingerprints, indicating their limited ability to cluster the data. In contrast, the MPNN embeddings showed some subtle clustering after t-SNE dimensionality reduction. Notably, the representations from the GMFU module exhibited significantly more pronounced clustering following t-SNE, suggesting an enhanced capacity to differentiate molecular efficacy in the high-dimensional space.

## 3 Discussion and Conclusion

In this work, we have proposed ChemXTree, a novel tree-enhanced graph representation model for binary classification tasks in small molecule drug discovery. The overall architecture first extracts initial molecular representations using graph neural networks. It then introduces a new module called GMFU for optimized feature selection. Finally, differentiable soft decision trees are employed as the output layer to enhance the model’s classification capability. We comprehensively evaluated ChemXTree on both the MoleculeNet benchmark and additional drug discovery datasets. The results demonstrated state-of-the-art performance, with ChemXTree outperforming current top models on 8 out of the 12 datasets. ChemXTree exhibited particular efficacy on tasks with limited samples. Furthermore, substitution experiments and ablation studies verified that connecting our proposed GMFU+Tree architecture after the graph encoding layer surpassed traditional FFN and XGBoost output layers. t-SNE results also indicated that the GMFU module contributes performance gains through feature selection. In summary, ChemXTree offers a new competitive end-to-end modeling scheme for drug discovery.

While effective on small datasets, ChemXTree’s performance on larger datasets needs improvement. Incorporating SOTA pre-trained GNN model could potentially improve ChemXTree’s scalability by leveraging more transferable features. Our future work will focus on GNN pre-training and feature extraction and balance input complexity with computational requirements. In addition, further simplifying the workflow and reducing the number of parameters are also key directions that need optimization for this framework. In conclusion, ChemXTree provides an effective solution for small sample learning, but expanding applicability to big data scenarios will require advancements in domains like pre-training, feature extraction, and workflow optimization. These important directions will form the core focuses of future research and development.

## 4 Materials and Methods

### 4.1 Datasets

MoleculeNet includes 16 datasets covering various scientific domains like Quantum Mechanics, Physical Chemistry, and Biophysics. These datasets are utilized for benchmarking machine learning models in the respective fields. Among the 16 datasets, there is several binary classification tasks in our comparison:

1. BBBP: The Blood–brain barrier penetration (BBBP) dataset. BBBP dataset evaluates the permeability of small molecules across the blood-brain barrier. It comprises 2039 validated compounds, featuring a positive-to-negative sample ratio of approximately 3.2:1.
2. BACE: The BACE dataset provides a comprehensive array of both quantitative (IC50 values) and qualitative (binary classification labels) data, evaluating the binding efficacy of inhibitors aimed at human *β*-secretase 1 (BACE-1). The dataset comprises 1,513 compounds, with an inactive-to-active label ratio of approximately 1.23:1.

- HIV: The HIV dataset originates from the Drug Therapeutics Program’s AIDS Antiviral Screen. It contains data on 41,127 compounds and their efficacy in inhibiting HIV replication. The dataset features a ratio of inactive to active compounds of approximately 27.5:1.
- ClinTox: The ClinTox dataset provides a comparative analysis between FDA-approved drugs and compounds that failed in clinical trials due to toxicity concerns. It includes 1,478 compounds, with ratios of FDA-approved to unapproved drugs at approximately 14.9:1, and clinical toxicity-positive to -negative compounds at approximately 12.2:1

For benchmarking on additional 8 ADMET datasets. Our ADMET datasets are derived from previous works. ^5,40^We thank Therapeutics Data Commons for providing access to these valuable resources.

Ames Mutagenicity (Toxicity): The Ames Mutagenicity dataset focuses on the potential mutagenic effects of compounds using the Ames test, which identifies substances that may damage DNA. ^44^ In this work, we refer to the Ames Mutagenicity dataset as AMES.

CYP2C9, CYP2D6, CYP3A4 Substrate(Metabolism): These datasets examine the roles of CYP450 enzymes 2C9, 2D6 and 3A4 in metabolizing endogenous and foreign compounds. The tasks of these datasets are predicting whether compounds act as substrates for these enzymes. ^45^

CYP2C9, CYP2D6, CYP3A4 Inhibition (Metabolism): In contrast to the Substrate datasets, these datasets primarily aim to predict whether compounds inhibit the metabolic activity of CYP450 enzymes 2C9, 2D6, and 3A4. ^**?**^

Bioavailability(Absorption): This dataset represents the rate and extent at which active molecules are absorbed from a drug product and become effective at the site of action. The dataset is used for predicting the activity of bioavailability. ^46^

### 4.2 Spilt and Evaluation Metric

Scaffold splitting is commonly adopted in cheminformatics as a way to better evaluate the model generalization. While random splitting offers simplicity in implementation, it often inadequately represents the model’s ability to generalize to novel data. Scaffold splitting, on the other hand, aims to divide the dataset into subsets based on structurally distinct molecules. This approach poses a greater challenge to learning algorithms compared to random splitting, thereby providing a more robust measure of generalization.

In the case of datasets from MolecuNet, we follow a previously established 8:1:1 ratio for splitting the data into training sets, validation sets, and test sets. For datasets that are not part of MoleculeNet, we employ a 7:1:2 saffold spliting. To establish a unified standard, we adhere to the guidelines set by MoleculeNet and other previous work, using ROC-AUC as the evaluation metric across all our experiments. This allowed direct comparison to existing benchmarks and studies. Adhering to MoleculeNet’s criteria, PRC-AUC is employed for datasets with a positive sample rate below 2%; otherwise, ROC-AUC is preferred. Therefore, we use ROC-AUC for evaluation in these datasets.

### 4.3 Comparison Models

N-Gram: N-Gram model introduced by Liu et al. offers a simple, unsupervised approach to represent molecules by embedding vertices and assembling them in short walks within the graph.

MolCLR: MolCLR proposed by Wang et al. is a framework for molecular representation learning that applies contrastive learning to encode molecular structures to potentially capture the underlying patterns and relationships in molecular data.

GraphMVP(Graph Multi-view Pretraining): GraphMVP developed by by Liu et al. represents a pretraining method for graph neural networks, leveraging self-supervised learning to exploit the relationships and consistencies between 2D topological structures and 3D geometric views.

PretrainGNN: PretrainGNN is a new strategy developed by Hu et al. for pre-training Graph Neural Networks (GNNs) that simultaneously trains on individual nodes and entire graphs, enhancing both local and global representations.

GEM(Geometry-enhanced Molecular representation): GEM improves molecular property prediction by integrating molecular geometry into a Graph Neural Network (GNN). It utilizes a specialized GeoGNN architecture to simultaneously consider the influence of atoms, bonds, and bond angles, thereby crafting a more detailed representation of molecules.

AttentionFP: AttentionFP is a model proposed by Xiong et al. This model employs a graph attention mechanism to focus on the most crucial parts of the molecular structure, particularly the nonlocal intramolecular interactions.

GAT (Graph Attention Networks): GAT is a class of graph neural networks distinguished by their utilization of attention mechanisms enabling GATs to selectively prioritize and integrate information from adjacent nodes.

MAT (Molecular Attention Transformer): MAT developed by Maziarka et al.is based on the Transformer designed for molecular representation. It enhances the self-attention mechanisms of the Transformer by incorporating inter-atomic distances and molecular graph structures, allowing for a deeper understanding of molecular features.

GROVER: This is a pretrained model designed by Rong et al. employs a self-supervised message passing transformer architecture, integrating message passing networks with a transformer framework to effectively learn molecular representations from unlabeled data. GROVER has two sizes: GROVERbase and GROVERlarge with different hidden layers.

InfoGraph: InfoGraph is an graph representation learning method proposed by Sun et al. It maximizes mutual information between graph-level and substructure representations, enabling robust graph-level learning.

Automated Machine Learning(AutoML): AutoML, denotes the process automation for selecting, finetuning, and training machine learning models. In this work, the input is sourced from a 2048-bit Morgan Fingerprints, followed by the automatic fine-tuning of a Feedforward Neural Network (FFN).

GraphFormer: GraphFormer is a novel architecture for molecular property prediction proposed by Yang et al. in 2021. It combines graph neural networks (GNNs) with the Transformer structure widely used in natural language processing.

Uni-Mol: Zhou et al. have developed Uni-Mol, a 3D molecular representation learning framework that incorporates a comprehensively pretrained model, adept at processing a broad spectrum of molecular and protein pocket data for diverse molecular representation tasks.

Among these models, N-gram, PretrainGNN, GROVER, GraphMVP, MolCLR, GEM and Uni-Mol are pre-train model and others are designed without pre-trained.

### 4.4 Message Passing Neural Networks Module

Our Message Passing Neural Networks(MPNN) Module is developed from the Chemprop framework and has undergone fine-tuning. To be more specific, as with many existing works on molecular graph representation, the initial required input is canonical SMILES without atom mapping. In our MPNN module, atoms and bonds of the molecules correspond to vertices V and edges E in graph structure, respectively. We encode the features of the atoms (atomic numbers, numbers of bonds, formal charges, chirality, quantity of hydrogen, hybridization, and aromaticity of the atom, atomic mass) and the features of the bonds (types, in conjugation/ring, stereochemical information) to obtain the initial vectors. Therefore we can get

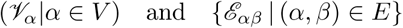

Furthermore, the initial directed edge features are obtained by connecting the atomic features of the first atom of the bond to the corresponding undirected bond feature ℰ_*α,β*_.Here, *ϕ*() is a simple function of concatenation.

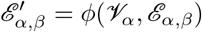

Subsequently, the initialized features are passed into the MPNN. Herein, the edge features are firstly processed through a simple network layer endowed with a ReLU activation function and learnable weights *W*_*i*_, where 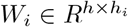. In our model, the values 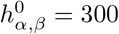 and 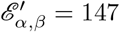 are taken. Therefore:

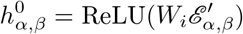

The directed edge attributes are subsequently updated through three iterative rounds of message passing, based on the local topology:

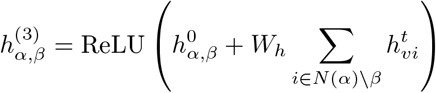

where *W*_*h*_ ∈ *R*^*h×h*^ and 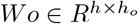
Let us denote

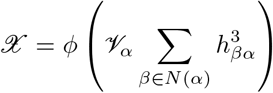

With this definition, the final hidden states can be expressed, which are then aggregated to form the atomic embeddings, as follows:

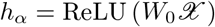

Therefore, the complete atomic embedding can be represented as:

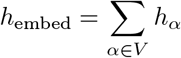

In this work, we do not present a new representational learning method.

### 4.5 GMFU Module

The attention component in the GMFU operates as follows:

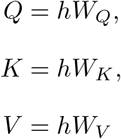

where *W*_*Q*_, *W*_*K*_, *W*_*V*_ ∈ *R*^*F ×d*^, *F* is the dimension of input features, and *d* is the dimension for each attention head.

*Q, K, V* are split along the last dimension *d* into *n*_heads_ different heads.

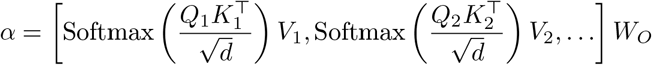

where, 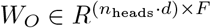.

After application of the self-attention mechanism, the feature vectors are pass into a gate control mechanism that plays a crucial role in the process of feature selection. This gate control mechanism has been adapted from GRU and GANDALF.

Inspired by previous works that have effectively utilized sparsity in neural networks, this work also adopts the t-softmax activation function proposed by Bałazy et al. in 2023 for feature masking. Specifically, the mask *M*_*n*_ is generated using the t-softmax function applied to a learnable parameter vector *F*_*n*_ and a tuning parameter *t*, as shown in the equation *M*_*n*_ = t-softmax(*F*_*n*_, *t*). This mask is then used to perform an element-wise multiplication with the input features *X*, resulting in the masked features *δ*_*t*_ as described in the equation *δ*_*t*_ = *M*_*n*_ *⊙ X*.

Similar to the standard GRU and GANDALF, we set the corresponding reset gate *r*_*t*_ and update gate *z*_*t*_ as follows:

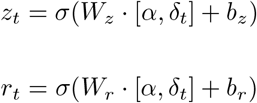

*σ* is the sigmoid function, which ensures the output is in the range of [0, 1]. With these gates, we can further specify the hidden state *H*_*t*_ and the candidate feature 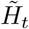 They are defined as:

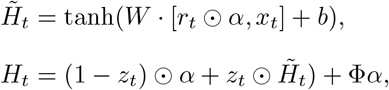

Here, Φ is a parameter of [0, 1]. The default value of Φ in the model is 0.05.

### 4.6 Differentiable Decision Binary Tree

We input the features processed through the GMFU into a differentiable binary decision tree. For such a decision tree, its role is to transform a *k*-dimensional input into a 2*D*-dimensional output. Although traditional decision trees have advantages like ease of deployment and straightforward algorithms, their methods are non-differentiable. Therefore, definite soft decision binary tree are considered in this work.

Inspired by prior research, ^29,47^ the *Soft Binning Function g* and the *decision stump o*_*i*_ in this work serve as specialized counterparts to the *splitting criterion* and the *decision node* found in traditional decision trees, respectively.

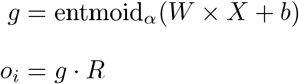

Where *W* ∈ *R*^*d×*2^, *b* ∈ *R*^*d×*2^

Definite decision binary trees in the model utilize all available features for each split through a linear combination of non-linear functions. A learnable feature mask *M* ∈ *R*^*d*^*′* is introduced to efficiently combine the outputs *o*_*i*_, allowing for scalability and comprehensive feature consideration.

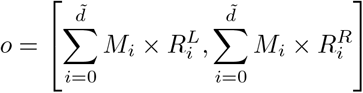

### 4.7 Training

The ChemXTree and baseline models were trained on the NYU Greene and NYUAD Jubail high-performance computing (HPC) clusters. Specifically, we utilized NVIDIA’s A100 80G GPUs for our experiments in most small datasets. For large size datasets, we escalated our computational capacity by deploying dual A100 80G GPUs. In every experiment conducted, drawing inspiration from the parameter tuning practices customary in traditional machine learning, we harnessed the prowess of Bayesian optimization. This method was utilized to meticulously traverse the parameter space comprising the learning rate, batch size, tree depth, tree breadth, the count of heads in the multi-head attention mechanism within GMFU, and the quantity of GRUs, alongside the dropout rate. This meticulous exploration was performed through 100 Bayesian searches in each experiment. In the ablation experiment and comparative experiment in ADMET datasets, we fine-tuned key parameters like learning rate, epoch, and batch size for each model. For the detailed hyperparameters, please see the Supplementary data.

## 5 Availability

All the data can be found in the https://moleculenet.org/ and https://tdcommons.ai/. The code for the models and results can be found in the GitHub repository accompanying this manuscript. The code for models and results will be available in the corresponding GitHub repository after this manuscript is published.

## 6 Acknoledgement

This work was supported by National Natural Science Foundation of China (Grant nos. 21933010, 22250710136, 22333006). We thank Ms.Bihui Guo of the University of Southampton for valuable discussions and contributions to the figures of our manuscript. We sincerely thank the High-Performance Computing (HPC) resources at NYU Abu Dhabi and New York University, as well as the dedicated staff and their technical support.

## 7 Supplementary data

**Figure S1:**
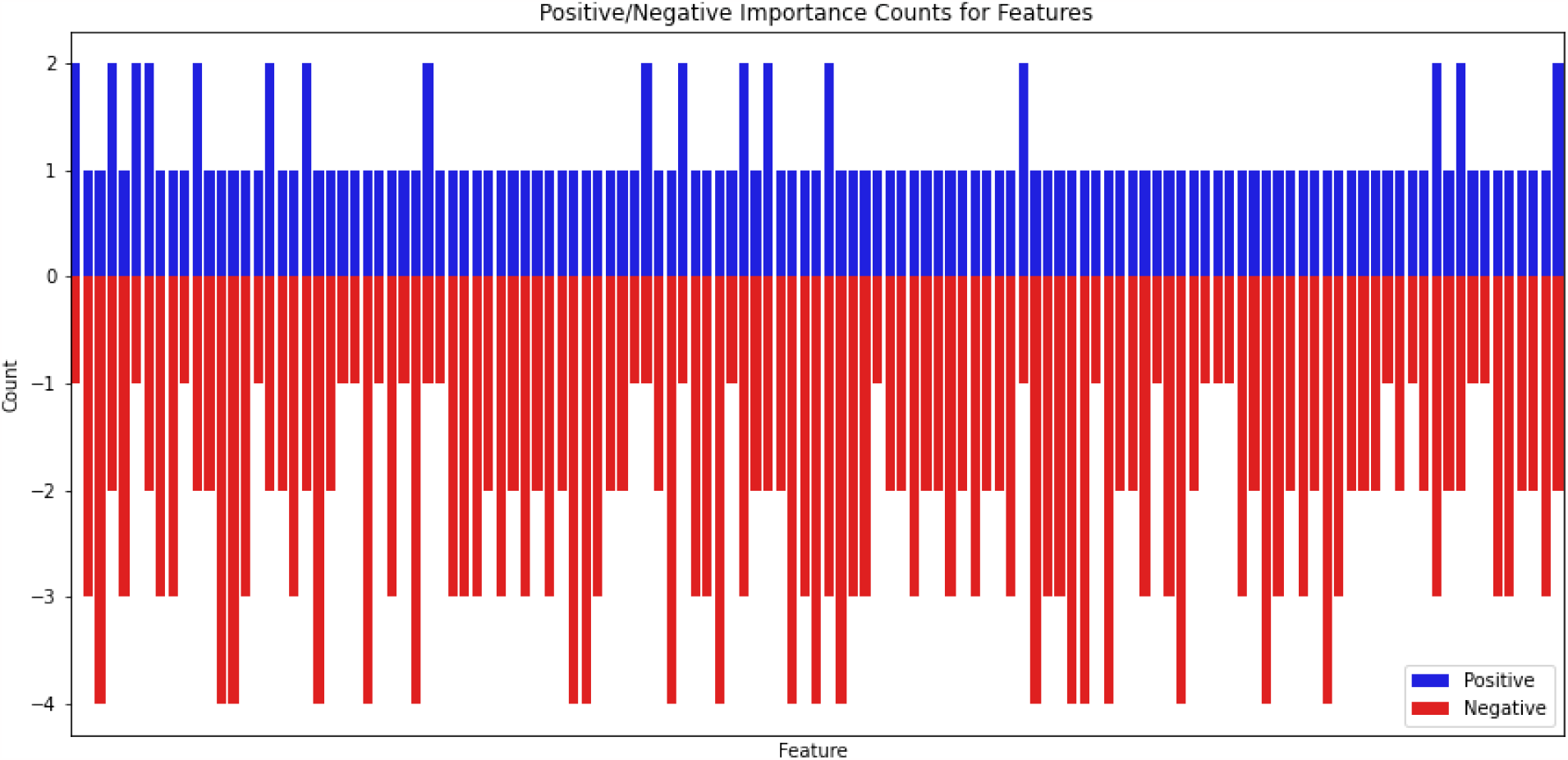
Part of the feature importance as determined by permutation experiments. Blue bars depict features that decrease model performance while red bars illustrate features increase performance, judged by ROC-AUC changes when removed. This result supports that employing a 5 independent training combined encoding helping the model better recognize important features

**Table S1:**
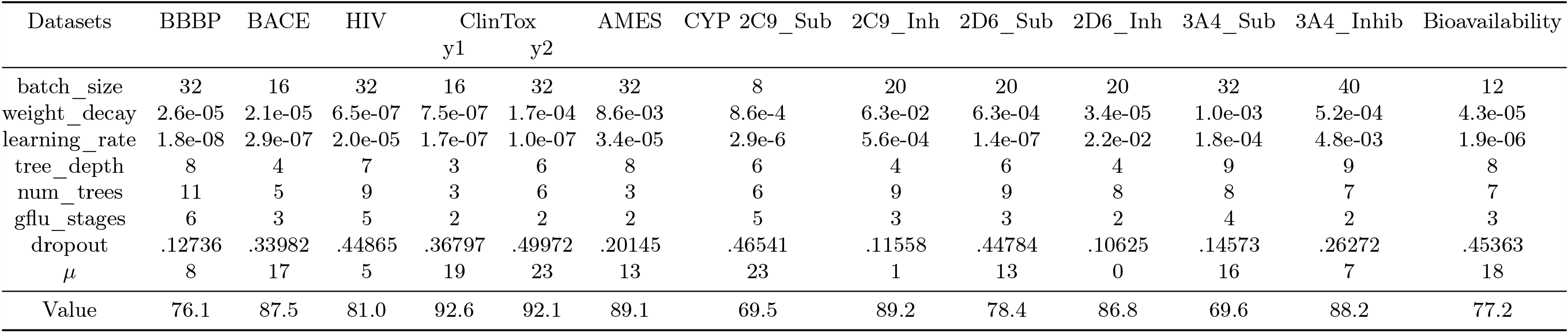
Parameter Information.

**Figure S2:**
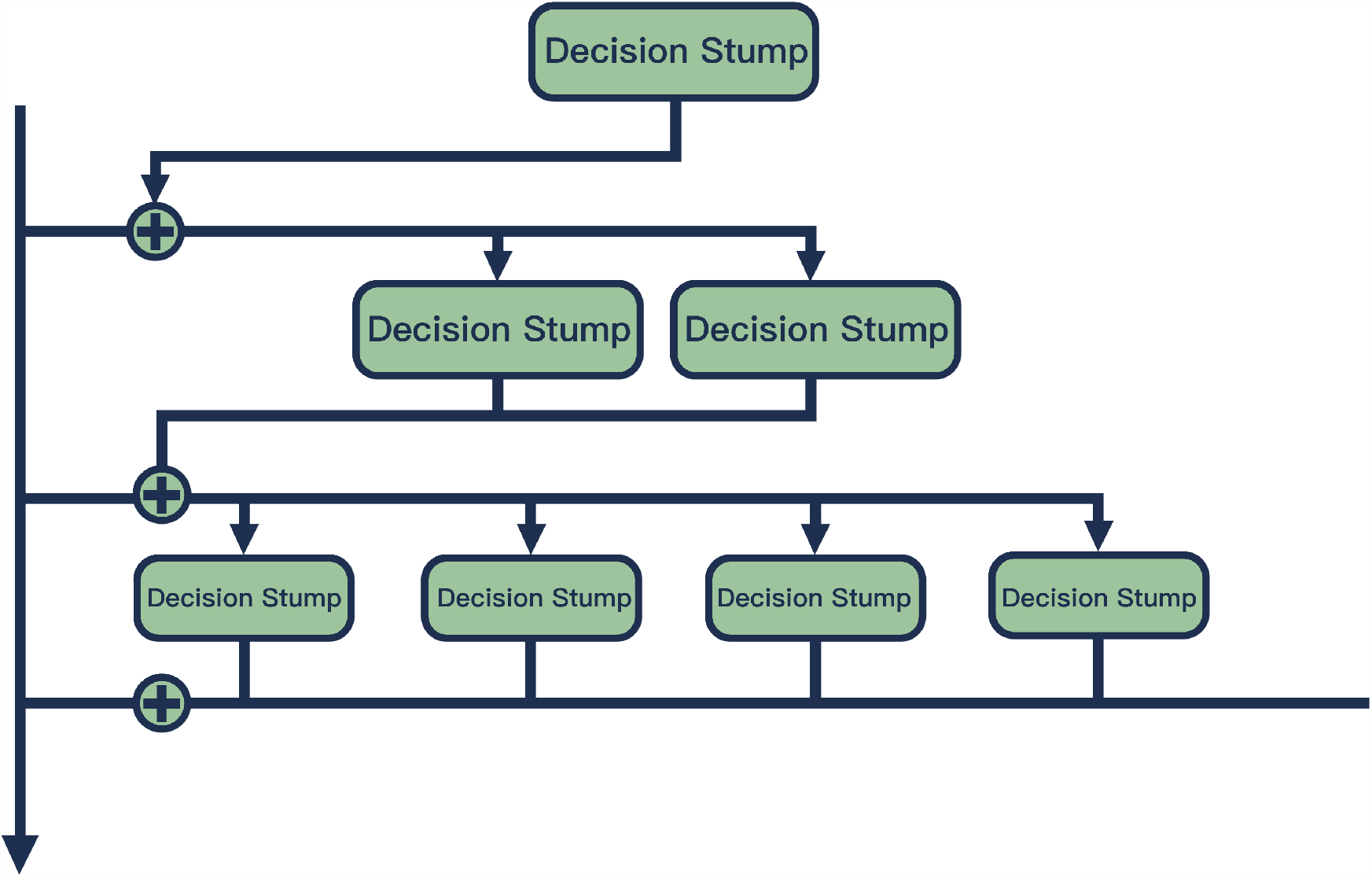
The framework of differentiable decision binary tree.

